# Engineering multi-degrading bacterial communities to bioremediate soils contaminated with pesticides residues

**DOI:** 10.1101/2024.02.13.580075

**Authors:** Sylvia Thieffry, Julie Aubert, Marion Devers-Lamrani, Fabrice Martin-Laurent, Sana Romdhane, Nadine Rouard, Mathieu Siol, Aymé Spor

**Affiliations:** INRAE, Institut Agro, Université de Bourgogne, Université de Bourgogne Franche-Comté, Agroécologie, Dijon, France; Université Paris-Saclay, AgroParisTech, INRAE, UMR MIA Paris-Saclay, 921120, Palaiseau, France

**Keywords:** bioremediation, ecological engineering, coalescence, glyphosate, isoproturon

## Abstract

Parallel to the important use of pesticides in conventional agriculture there is a growing interest for green technologies to clear contaminated soil from pesticides and their degradation products. One specific technique, the inoculation of degrading micro-organisms in polluted soil, known as bioaugmentation, is a promising method still in needs of further developments. Specifically, improvements in the understanding of how degrading microorganisms must overcome abiotic filters and interact with the autochthonous microbial communities are needed in order to efficiently design bioremediation strategies. Here we designed a protocol aiming at studying the degradation of two herbicides, glyphosate (GLY) and isoproturon (IPU), *via* experimental modifications of two source bacterial communities. We used statistical methods stemming from genomic prediction to link community composition to herbicides degradation potentials. Our approach proved to be efficient with correlation estimates over 0.8 - between model predictions and measured pesticide degradation values. OTUs significantly associated with the degradation ability, and therefore identified as relevant by the models were confronted to the literature. Next, multi-degrading bacterial communities were obtained by coalescing bacterial communities with high GLY or IPU degradation ability based on their community-level properties. Finally, we evaluated the efficiency of constructed multi-degrading communities to remove pesticide contamination in a different soil. While results are less clear in the case of GLY, we showed an efficient transfer of degrading capacities towards the receiving soil even at relatively low inoculation levels in the case of IPU. Altogether, we developed an innovative protocol for building multi-degrading simplified bacterial communities with the help of genomic prediction tools and coalescence, and proved their efficiency in a contaminated soil.

## Introduction

Pesticides are still widely used in conventional agriculture to maintain high yield by controlling pests among which broadleaf weeds competing with the crops (1). Yet their persistence in soil and their transfer to water resources cause ecological damages that are increasingly documented (2). Indeed, pesticides or their degradation metabolites can have negative effects on animals and microorganisms which in turns affect ecosystem functions and services (3; 4).

Recurrent pesticides exposure can lead to the selection of degrading populations among soil bacterial community that are able to use pesticides as nutrient and energy source for their growth (5). Pesticide biodegradation by soil microorganisms has been amply documented and represents a precious baseline to develop bioremediation strategies of polluted ecosystems (6; 7). Classically, bioaugmentation strategies relies on the isolation of specific degrading strain(s) by enrichment in a selective medium supplemented with the pesticide as sole carbon or nitrogen source. This process is time consuming and assumes that the degrading bacteria can survive and grow in laboratory conditions, which is known to be the case for only a small fraction of all microorganisms (8). Besides isolation, multiple studies showed that bioaugmentation with pure degrading strains or simple bacterial consortia turned out to be inefficient in field-scale experiments, due to establishment and survival problems of microbial inocula or to an *in situ* decline of their degrading activities (9; 10).

In order to overcome these limitations, we propose to adopt the vision of quantitative geneticists who are trying to introgress a genomic region into a given genetic background to select a phenotype of interest. By doing so, pesticide-degrading bacteria are seen as the alleles responsible for the phenotype of interest *i. e.* pesticide biodegradation, that need to be introgressed into a contaminated soil microbial community, corresponding to the genetic background to improve. Hence, the microbial community matches the plant or animal individual as the experimental unit. Therefore, we transposed tools from genomic selection used by quantitative geneticists to microbial ecology with a bioremediation perspective.

Statistical methods used in genomic selection result from a mix of two historical approaches: the Best Linear Unbiase Predictor approach (BLUP), blind to genetic trait determinism, uses kinship level between individuals to derive the proportion of expected shared alleles and guides choice for individuals to cross (11; 12). The second approach, akin to the QTL approach is based on genomic markers to identify regions implied in the expression of phenotypes of interest (13). The emergence of NGS (Next Generation Sequencing) opened the possibility to precisely estimate shared alleles between individuals, allowing the use of BLUP type models based on the realised, instead of expected, kinship matrix. We can then use individual genotype as a proxy to predict the genetic value. Correspondingly, one could develop a prediction model for microbial community “phenotype” (community function) from their “genotype”, which consists of its specific composition determined by sequencing of taxonomic markers (*e.g.* 16S rDNA amplicon).

To test the relevance of this original approach, we focused our research on two herbicides: Glyphosate (GLY) and Isoproturon (IPU) and experimentally obtained a cohort of microbial communities degrading these two xenobiotics. After extraction of degrading communities, one for each of the two herbicides, serial dilutions coupled with a range of biocidal treatments were performed in order to obtain compositional variants. Importantly, they were inoculated in the recipient soil previously sterilized, as a mean to select degrading bacteria able to grow in this specific environmental niche (14). Indeed, environmental filtering by pH, soil structure, organic matter proportion, is thought to be a major cause for the inefficiency of strain bioaugmentation (15). For each of the compositional variant, degrading capacity was measured by radiorespirometry and 16S rDNA sequencing were undertaken, allowing us to explore the links between the pesticide-degrading function and the bacterial community composition at the Operational Taxonomic Unit (OTU) level. We used statistical methods routinely used in genomic prediction, focusing on two models derived from linear regression, Ridge Regression (RR) and LASSO, as well as a nonlinear machine-learning method: Random Forest (RF) (16). We then investigated the explanatory power of the models and looked at highlighted OTUs. To go further, coalescence of GLY and IPU degrading communities, specifically chosen based on their intrinsic properties such as diversity levels, degrading capacities and relative abundance of OTUs of interest, was performed in order to explore the compatibility of degrading communities. The multi-degrading communities were finally inoculated in a recipient inhabited soil to evaluate the efficiency of the proposed protocol to bioremediate a pesticide contaminated soil.

## Materials and methods

### Soil sampling

We collected three different soils based on their potential to degrade two herbicides, glyphosate and isoproturon. One soil was collected from Noiron-sous-Gevrey (France, 47°11’36.38”N, 5°4’55.44”E) for its potential to degrade glyphosate (GLY). It contains 42% silt, 42% clay and 15% sand and shows a pH of 8.04, as well as 25.37 g/kg organic Carbon and 2.15 g/kg Nitrogen. A second was collected from the Epoisses INRAE experimental farm (Bretenière, France, 47°30’22.18”N, 4°10’26.46”E) for its potential to degrade isoproturon (IPU) with soil properties as follows: 51.9% silt, 41% clay, and 6.2% sand, pH 7.2, and organic Carbon and Nitrogen content 15.5 g/kg and 1.4 g/kg dry soil, respectively. A third soil was collected from the CEREEP Ecotron Research Station (Saint-Pierre-lès-Nemours, France, geographic coordinates: 48°16’58.9”N, 2°40’19”E) and displayed a poor degradation potential for both herbicides. This last soil is a cambisol with moor composed of 19% silt, 6.98% clay, 74.1% sand, and with 14.7g/kg organic C; 1.19g/kg N; pH of 5.22. All three soils were sieved at 4mm, and a sufficient amount of the Ecotron soil was sterilised by *γ* sterilisation (2 times 35 kGy in Conservatome, Dagneux, France). The Noiron-sous-Gevrey and Epoisses soils were pre-incubated for 30 days in 300 g microcosms in triplicates with their attributed herbicide (GLY and IPU, respectively at days 1 and 15 according to their agronomical dose, *i.e.* 3 mg/kg equivalent dry mass (edm) for GLY and 2mg/kg edm for IPU).

### Glyphosate and Isoproturon mineralisation potentials

Mineralisation ability of the two herbicides were measured on 15 g edm soil samples that were adjusted to the same moisture level (70% of water holding capacity (WHC) of the soil). Solutions of C-ring labelled IPU and C-GLY (on C-P bond) were obtained at a concentration reaching 1 667 Bq radioactivity per dose (400*µ*l of herbicide solution added to the samples at agronomical dose). All samples were then placed in closed jars along with 10ml water in an open container to maintain humidity and 5ml sodium hydroxide NaOH at 0,2M to trap emitted CO_2_. The jars were put in the dark at room temperature (20°C) for 42 days. The NaOH containers were regularly replaced for temporal assessments of emitted CO_2_ with addition of 10ml scintillation fluid (ACSII, Amersham) and measures on a Beckman scintillation counter. For each herbicide, the mineralisation potential was expressed as the cumulative percentage of CO_2_ evolved from the initially added C-pesticide over time.

### Microbial community variants construction

For each herbicide-degrading soil microbial community, compositional variants were obtained by coupling serial dilutions with biocidal treatments, as described in Romdhane et al., 2021 (17). Each soil community was first extracted by blending 33 g edm of soil with 60 ml sterile distilled water, then \diluted 10 times with sterile 0.9% NaCl solution to get the 10 dilutions. 10^-1^, 10^-2^ and 10^-3^ dilutions were retrieved by 10 times serial-dilutions. For each dilution level, we applied 9 different biocidal treatments (see Table 1 for more details) in triplicates and kept untreated samples as controls. This experimental procedure led to a total of 120 community suspensions (= 4 dilutions x 10 treatments x 3 replicates) per herbicide-degrading community. A volume of 10.5ml from each suspension was inoculated into 50g dry sterilized Ecotron soil in plasma flasks closed with sterile cotton lids, with addition of 2ml of the adequate herbicide solution to reach the agronomical dose in order to facilitate the establishment of the degrading community. All microcosms were kept for 43 days in the dark at room temperature (20°C) and moisture was adjusted to vary between 50% to 70% of WHC by addition of sterile water. After 43 days, 250mg edm of soil was collected for each microcosm to assess microbial community composition using 16S rDNA sequencing, 5g to determine relative humidity and 15g to assess herbicide mineralisation potentials.

### Assessment of microbial community composition and diversity

DNA was extracted from 250mg edm for the 246 samples (compositional variants and original soils) using the DNeasy PowerSoil-htp 96 well DNA isolation kit (Qiagen, Hilden, Germany). Generation of amplicons for Illumina MiSeq sequencing was done according to a two steps PCR protocol as follows: *i*) amplification in duplicates of the V3-V4 hyper-variable region of the bacterial 16s rRNA with fusing primers U341F (5’-CCTACGGGRSGCAGCAG-3’) and 805R (5’-GACTACCAGGGTATCTAAT-3’), and, to allow the subsequent addition of multiplexing index-sequences, overhang adapters (forward: TCGTCGGCAGCGTCAGATGTGTATAAGAGACAG, adapter: GTCTCGTGGGCTCGGAGATGTGTATAAGAGACAG). PCR cycles started at 98°C for 30s, then 55°C for 30s and 72°C for 30s and a final extension for 10min at 72°C, *ii*) pooling of duplicates and amplification with unique pairs of tag per sample, at PCR cycles starting with 8°C for 3min followed by 98°C for 30s, 55°C for 30s, 72°C for 30s and a final extension for 10min at 72°C. 2% agarose gel were prepared to visually verify amplicon sizes, and we noted two samples (both from the GLY compositional variants) without amplicons that were taken out for downstream analyses. All samples were then cleaned with SequalPrep Normalization plate kit 96-well (Invitrogen, Carlsbad, CA, USA), pooled and sequenced on MiSeq (Illumina, 2 x 250 bp) using the MiSeq reagent kit v2 (500 cycles).

The sequence data were analysed using an in house developed Jupyter Notebook (18) streaming together different bioinformatics tools. Briefly, R1 and R2 sequences were assembled using PEAR (19) with default settings. Further quality checks were conducted using the QIIME pipeline (20) and short sequences were removed (< 400bp). Reference-based and *de novo* chimera detection, as well as clustering in OTUs were performed using VSEARCH (21) and the adequate reference databases (SILVA’s representative set of sequences). The identity thresholds were set at 94% based on replicate sequencing of a bacterial mock community containing 40 bacterial species. Representative sequences for each OTU were aligned using MAFFT (22) and a 16S phylogenetic tree was constructed using FastTree (23). Taxonomy was assigned using BLAST (24) and the SILVA reference database v138 (ref 25). Diversity metrics, that is, Faith’s Phylogenetic Diversity (PD) (26), richness (observed species) and evenness (Simpson’s reciprocal index), describing the structure of microbial communities were calculated based on rarefied OTU tables (10 000 sequences per sample). UniFrac distance matrices (27) were also computed to detect global variations in the composition of microbial communities. OTUs showing low abundance were discarded with a threshold of 210 counts across the compositional variants, to avoid artefactual association of rare OTUs with degradation potential and to lower the number of parameters (OTUs) to be estimated, from 5,145 to 1,316. In addition, one GLY compositional variant had a read depth below 5,000 and was then removed for downstream analyses (mean of count per community around 16,700).

### Co-occurrence networks and visualisation of OTU predictors

Bacterial co-occurrence networks were constructed based on filtered OTU count data (1316 OTUs) over the 240 samples degrading communities using a sparse multivariate Poisson log-normal (PLN) network model with a latent Gaussian layer and an observed Poisson layer using the R package PLNModels v0.11.04 (ref. 28; 29). The best model was selected using a Stability Approach to Regularisation Selection (30). Phylogenetic relationships between OTU predictors were visualized using the Interactive Tree of Life webservice (31).

### Predicting herbicide mineralisation potential from microbial community composition

We derived the principle of predicting herbicide mineralisation potential from the bacterial community composition from the same logic as used in genomic prediction, where a model is trained to predict phenotype on the basis of the genotype of individuals as assessed through numerous molecular markers. Here we trained models to predict herbicide degradation based on the composition of bacterial communities. All the following analyses are based on three statistical methods used in genomic prediction: Ridge Regression (RR), LASSO and Random Forest (RF). The first 2 methods build a regression model based on the relative composition in OTUs of the compositional variants over the function performed by their community. This leads to dealing with a high number of parameters to be estimated (the effect of each of the 12 1316 OTUs, namely) compared to the number of experimental samples (120 compositional variants per herbicide-degrading community). In this so-called large *p* small *n* setting, using a regular linear regression (Ordinary Least-Square) would lead to inflated variance of the estimates. To alleviate this problem, penalized regression methods use various penalty functions to shrink all estimates (Ridge Regression, RR) and/or perform variable selection (in the case of LASSO). Here we used a Bayesian implementation of these 2 penalized regression methods with prior distribution of bacterial species or OTU effects chosen as a Gaussian Distribution for RR and a Laplace distribution for LASSO. They were run using the R package BGLR v1.1.0 (ref. 32). The third method, Random Forest, is an ensemble method based on building decision trees to determine the importance of the OTUs by partitioning them depending on their counts in relation with the measured community function (33). This method uses a combination of tree predictors and has the advantage of not hypothesizing additive effects of the OTUs and can thus be powerful in non-additive situations. The R package RandomForest v4.7-1.1 (ref. 34) was used for the following analysis, with parameters: number of trees of 500, node size of 5 and tested variables at each split of 1000.

Prior to any analysis, OTUs counts were corrected for sequencing depth (data/community sum * max of all community sum) then centered and reduced. Based on the mineralisation curves (see Supp. Fig. 1), we estimated 3 measurable curve parameters that where further considered as our community phenotypes: the total cumulative mineralisation after 42 days (Total), mineralisation value at the time point at which the variance across samples of the cumulative mineralisation potential is maximised (HighVar) and the slope of the linear regression of the cumulative mineralisation potential over the first 10 days (Slope). This last parameter approximates the mineralisation potential rate. Note that for GLY, HighVar is estimated at day 42, and is therefore equivalent to Total. On the opposite, for IPU, HighVar landed on day 13. To assess the prediction potential of these 3 statistical methods, we used a cross-validation scheme using 4/5 of the data to train the models, and the remaining 1/5 of the data to test its accuracy. The process was repeated 8 times, redrawing the sets each time. The metric we used to assess the accuracy of prediction was the Pearson correlation between predicted and measured herbicide mineralisation potential.

### Mixing herbicide-degrading compositional variants

We created artificial microbial communities by mixing GLY-degrading with IPU-degrading compositional variants. The modalities used to create these communities were driven by different hypotheses leading to six scenarios based on community structure or degrading features: *i*) measured mineralisation rate for the herbicide they were selected for, *ii*) diversity of the compositional variants, *iii*) predicted mineralisation potential rate of the other herbicide and *iv*) abundance of OTUs detected as having positive or negative associations with the measured mineralisation rate. For each scenario, 5 IPU-degrading communities and 5 GLY-degrading communities were selected and mixed by random pairing. The 6 different scenarios were (Fig. 1):

- DivPhyl+: Compositional variants displaying both a high phylogenetic diversity (for GLY: selected communities mean of 94.6 on the overall mean of 56.8, for IPU: selected communities mean of 86 on the overall mean of 52.3) and a high mineralisation rate (in top 20% for IPU degrading communities and top 25% for GLY).
- DivPhyl-: Compositional variants displaying both a low phylogenetic diversity (for GLY: selected communities mean of 62.8 on the overall mean of 56.84, for IPU: selected communities mean of 45.8 on the overall mean of 52.3) and a high mineralisation rate (in top 20% for IPU degrading communities and top 25% for GLY).
- Predicted+: Compositional variants displaying both a high experimentally measured mineralisation rate for the herbicide they were selected for (top 10% for GLY and top 55% for IPU) and a high predicted mineralisation potential rate for the other herbicide (estimated as the slope by Bayesian LASSO model, selected communities in top 6% of IPU predictions on GLY communities and in top 35% of GLY predictions on IPU communities).
- Predicted-: Compositional variants displaying both a high experimentally measured mineralisation rate for the herbicide they were selected for (top 30% for GLY and top 55% for IPU) and a low predicted mineralisation potential rate for the other herbicide (estimated as the slope by Bayesian LASSO model, selected communities in bottom 40% of IPU predictions on GLY communities and in bottom 50% of GLY predictions on IPU communities).
- OTUMinMax: Compositional variants maximizing the abundance of OTUs detected as having significant and strong positive associations while minimizing the abundance of OTUs detected as having significant and strong negative associations with the measured mineralisation rate (based on the Slope parameter and estimated by Bayesian LASSO model).
- OTUMax: Compositional variants maximizing the product of the abundance of OTUs detected as having significant and strong positive associations with the measured mineralisation rate times their estimates (based on the Slope parameter and estimated by Bayesian LASSO model).

**Figure 1:**
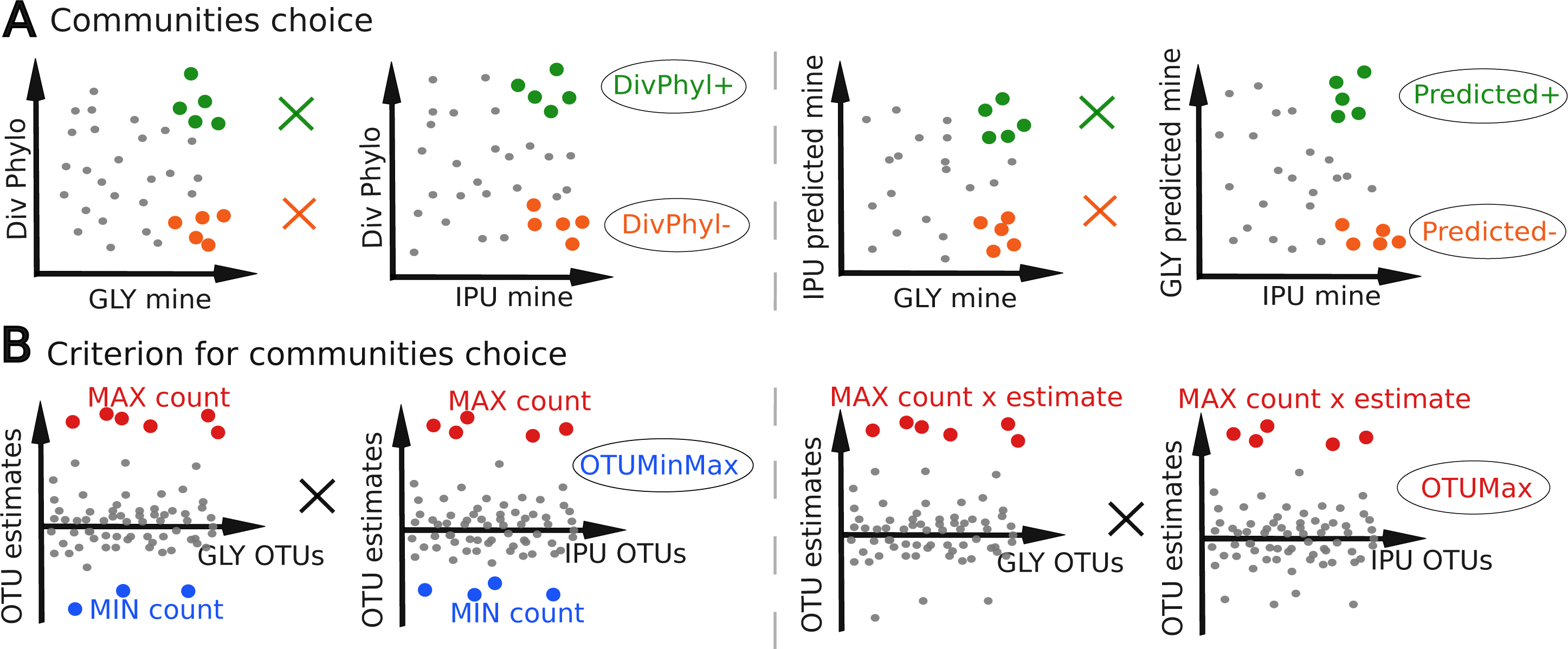
Graphical representation of criterion for selection of IPU and GLY communities to be mixed, under 6 scenarios. Crosses represent the mixing, always between a community originated from GLY degrading soil and one originated from IPU degrading soil. In panel A (4 scenarios), choice is made on communities’ characteristics: measured pesticide mineralisation rate (mine), phylogenetic diversity (Div Phylo) and predicted mineralisation, by BL model for Slope (predicted mine). Colored dots are the chosen communities to be mixed. In Panel B (2 scenarios), community choice is based on OTUs estimates (BL model for slope phenotype). Estimates represent the strength of the association between the abundance of a given OTU and the phenotype. Coloured dots are the OTUs taken in account for the choice, so OTUs displaying high estimates (in red) or low estimates (in blue). Scenario OTUMinMax maximizes the abundance of OTUs with high estimates and minimizes OTUs with low estimates while OTUMax maximizes the product of abundance of high estimates OTUs and their estimates.

For community mixing, 2.5g of each parental community was added in 50g sterilised dry Ecotron soil. All microcosms were then watered up to 70% of soil water holding capacity and boosted by the addition of an agronomical dose of both IPU and GLY, closed with sterile cotton lids and incubated for 45 days. The resulting 30 microcosms were then sampled, and the GLY and IPU mineralisation potential of the resulting microbial communities were then assessed using radiorespirometry.

### Coalescence of the herbicide multi-degrading communities into the original non-degrading soil

We selected the 5 best communities derived from the mixing of compositional variants based on their efficient IPU mineralisation and their improved GLY mineralisation compared to their parents. 4 levels of inoculations were prepared for each of these communities, reaching 50 g edm of soil in each microcosm. Inoculations were handled by mixing 0.1 g, 0.5 g, 1 g or 5 g (corresponding respectively to 0.2%, 1%, 2% and 10% mixing-ratio) of soil of the best communities with the original, non-sterilised Ecotron soil, with addition of 2 ml of both herbicide solutions to reach the agronomical dose, in order to facilitate the establishment of the degrading communities and mimic an herbicide contamination. IPU and GLY mineralisation potential of the coalesced communities were then assessed by radiorespirometry.

## Results

### Evaluation of the GLY and IPU mineralisation potential of the three selected soils

Mineralisation potentials of IPU and GLY differed greatly between soils, with the Ecotron soil showing the lowest potential for both pesticides (Supp. Fig. 2). IPU is far better mineralised in Epoisses (27.3% ±1.4) soil as compared to the two other soils (6.5% ±0.04 for Ecotron and 14.7% ±0.2 for Noiron). The Noiron soil exhibits the highest GLY mineralization (30.0% ±0.6), although the difference with the other soils is less striking than it is with IPU (17.9% ±0.1 for Ecotron and 21.6% ±0.5 for Epoisses).

### GLY and IPU-degrading compositional variants

We coupled serial dilutions with different biocidal treatments to construct compositional variants of GLY-degrading bacterial communities from Noiron soil and IPU-degrading ones from Epoisses soil (Table 1). Principal coordinates analysis of the weighted Unifrac distance matrix revealed a continuous variation in composition across both IPU and GLY communities (Supp. Fig. 3). Compositional variants are significantly different from the original communities found in IPU and GLY degrading soils. This indicates that community manipulation was successful to construct compositional variants for both GLY and IPU original communities.

We then evaluated mineralisation potential for both GLY and IPU-degrading compositional variants. Concerning GLY communities, the overall temporal dynamics of GLY mineralisation curves were similar between the different treatments with a slow saturation yielding in a cumulative mineralisation after 42 days ranging between 12.3% and 40.4% of -IPU initially added to microcosms. Moreover, we found a strong and significant impact of soil suspension dilution on GLY mineralisation potential, with a marked decreased GLY mineralisation in the more diluted soil suspensions (Fig. 2 A & C). Specific treatments (HS, Ox1, pH11) displayed a stronger reduction of total mineralisation compared to the control microcosms (Fig. 2 C). Furthermore, we noted that none of the compositional variants reached the total mineralisation potential of the original Noiron soil. This points out an overall decline either due to the experimental manipulation of soil suspensions or to differences in physico-chemical properties of the receiving soil (sterilized Ecotron soil), both potentially affecting GLY mineralisation capacities.

**Figure 2:**
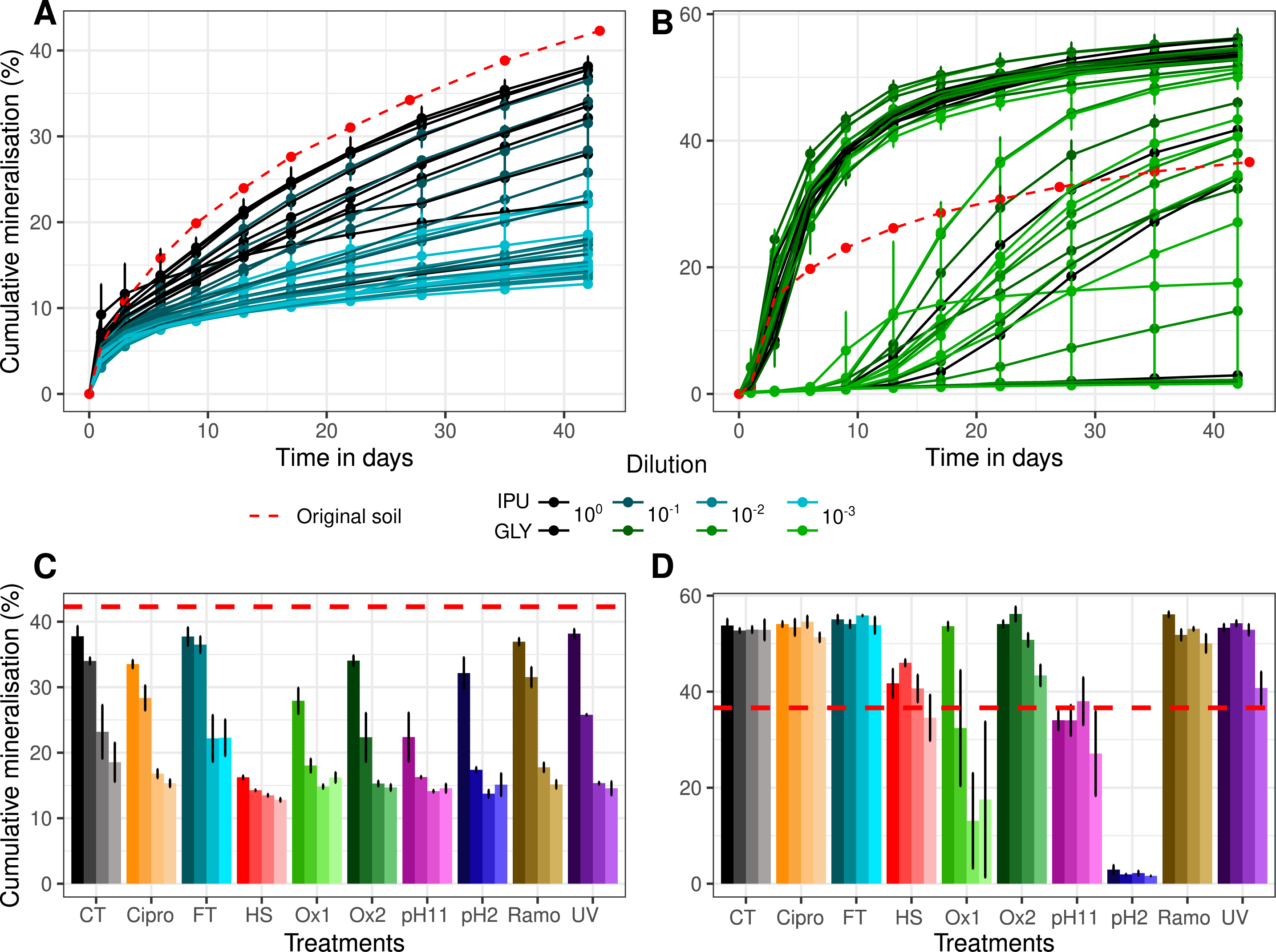
Kinetics of mineralisation and cumulative mineralisation of GLY and IPU after 42 days of incubation. Potential of GLY mineralization of communities derived from Noiron soil (Panel A & C) and of IPU mineralization from Epoisses soil (Panel B & D). Lines of bars correspond to one condition of treatments and dilutions with error bars as standard error of the mean (n = 3). Dashed lines are for original soil community, dotted and dashed for original soil communities after 30 days pesticide incubation. Biocidal treatments: Cipro = ciproflaxin (antibiotics), FT = Freeze-thaw, HS = Heat shock, Ox1 = strong oxidation, Ox2 = mild oxidation, pH11, pH2, Ramo = Ramoplanin (antibiotic) and UV = exposition to UV.

Regarding IPU communities, three distinct dynamics can be identified: *i*) some microcosms displayed IPU mineralisation kinetics with a rapid start reaching a plateau in about 15 days with a high final cumulative mineralisation (around 50% to 60%, even higher than the original Epoisses soil that only reached 36%), *ii*) some microcosms displayed kinetics with a lag phase of about 10 days followed by an increase of mineralisation rate up to reaching a plateau at the end of the experiment, with a total cumulative mineralisation ranging between 30% to 50% and *iii*) some microcosms displayed nearly no mineralisation with cumulative mineralisation after 42 days below 4% (Fig. 2 B & D). The acidic treatment (pH2) had the strongest negative impact on IPU mineralisation, followed by the basic (pH11) and heat shock (HS) treatments. Interestingly, diluting soil suspensions did not have an effect on IPU mineralisation as observed for GLY communities. However, for some treatments (*e.g.* for Ox1, Ox2 and pH11 treatments) dilutions increased the variance between triplicates.

### Association between bacterial community composition and herbicide mineralisation potential

The three aforementioned statistical methods (BRR, BL and RF) were applied to our set of community variants. Specifically, we used the normalized OTU table as the set of predictors, for which effects have to be estimated, and three parameters of herbicide mineralisation kinetics as the predicted variables: the Slope on the mineralisation curve over the first 10 days, the cumulative mineralisation (Total) and the percent of pesticides mineralisation at the time point where the variance is the highest (HighVar). Correlations between the actual and predicted variables after cross-validation depicted in figure 3 are overall very high, often with correlation coefficient well above 0.75. Unsurprisingly, correlations on training data are close to 1 (higher than 0.95 for IPU communities, and ranging from 0.89 to 0.98 for GLY) with very low variance. Looking at the prediction accuracy on testing sets, the values are also higher for IPU than for GLY, with the exception of the Total for which correlations went down from above 0.9 to 0.8-0.85. Prediction accuracy for GLY communities are comprised between 0.77 and 0.85 with no clear difference between the 3 predicted variables. We also did not detect any clear difference in the prediction accuracy of the different statistical methods. Even if RF yielded the highest correlation on the training set, its results on the testing set were in the same range as the two penalized regression methods.

**Figure 3:**
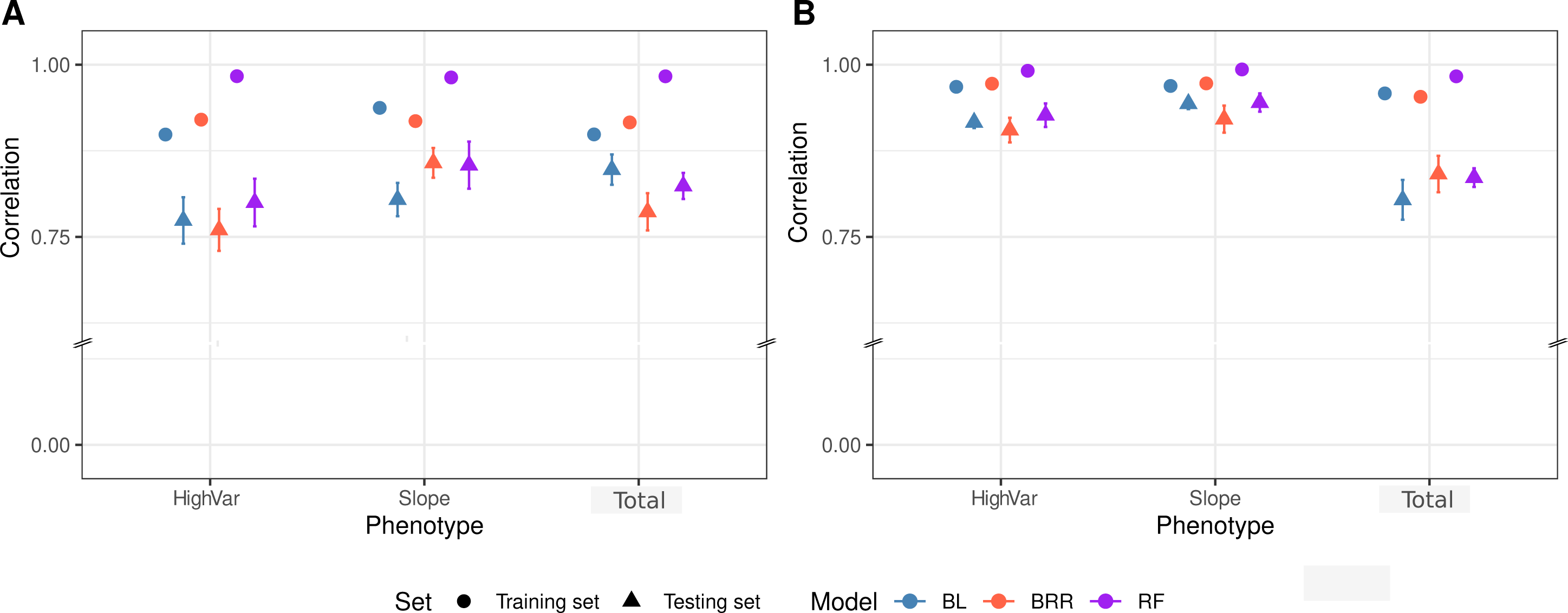
Prediction accuracy assessed by cross-validation on mineralisation data. Training set on 4/5 of the samples, testing set on the left 1/5, shuffled between the 8 runs (n=120 for each herbicides). Errors bars represent standard error of the mean for A) glyphosate mineralisation and B) isoproturon mineralisation. Phenotypes are mineralisation kinetic parameters as follows: HighVar the mineralisation value at the time point at which the variance across samples of the cumulative mineralisation potential is maximised, Slope the slope of the linear regression of the cumulative mineralisation potential over the first 10 days and Total the total cumulative mineralisation after 42 days.

The capacity of the different methods to accurately predict mineralisation parameters was further investigated by looking at OTUs with a strong effect on mineralisation, *i. e.* having a large estimated effect size, either positive or negative. Interestingly, detected OTUs by RF were overall largely different from the ones detected by BL and BRR for which 28 OTUs were common over the 30 most important detected for Slope variable for both pesticides (Supp. Fig. 4). We therefore looked in more details at the taxonomy and the partial correlation patterns between OTUs detected by both BL and BRR models applied on the Slope and Total (Fig. 4). Overall, there are less negative than positive estimates, and their amplitudes are lower than positive ones. In addition, for the IPU communities, OTUs that are detected as having an effect on the Slope variable were also detected as important for the Total (19 out of 27), which is not the case for the GLY communities (7 out of 26). Co-occurrence pattern (positive partial correlations) are found across the displayed OTUs and they are particularly present between taxonomically close OTUs *e.g.* across *Pseudomonas* genus or Cytophagales. Interestingly, a recurrent pattern of taxonomically-related OTUs showing opposite estimates was detected for both herbicides (*e.g.* Actinomycetales order in GLY and IPU, *Pseudomonas* genus, Sphingobacteriales and Bacilliales in IPU). Some of the selected OTUs are also important for degradation prediction by the RF method and even if they are not the ones with highest estimates (as determined by BL), they can be seen as robust because detected by all three methods. Moreover, 6 OTUs are shared by GLY and IPU linear prediction models, underlying either multi-degrading strains or more likely strains involved in the downstream part of mineralisation pathways. They could also be facilitators of degrading strains, altering the abiotic conditions favourably (or unfavourably for Rhizobiales *Balneimonas* in the case of IPU).

**Figure 4:**
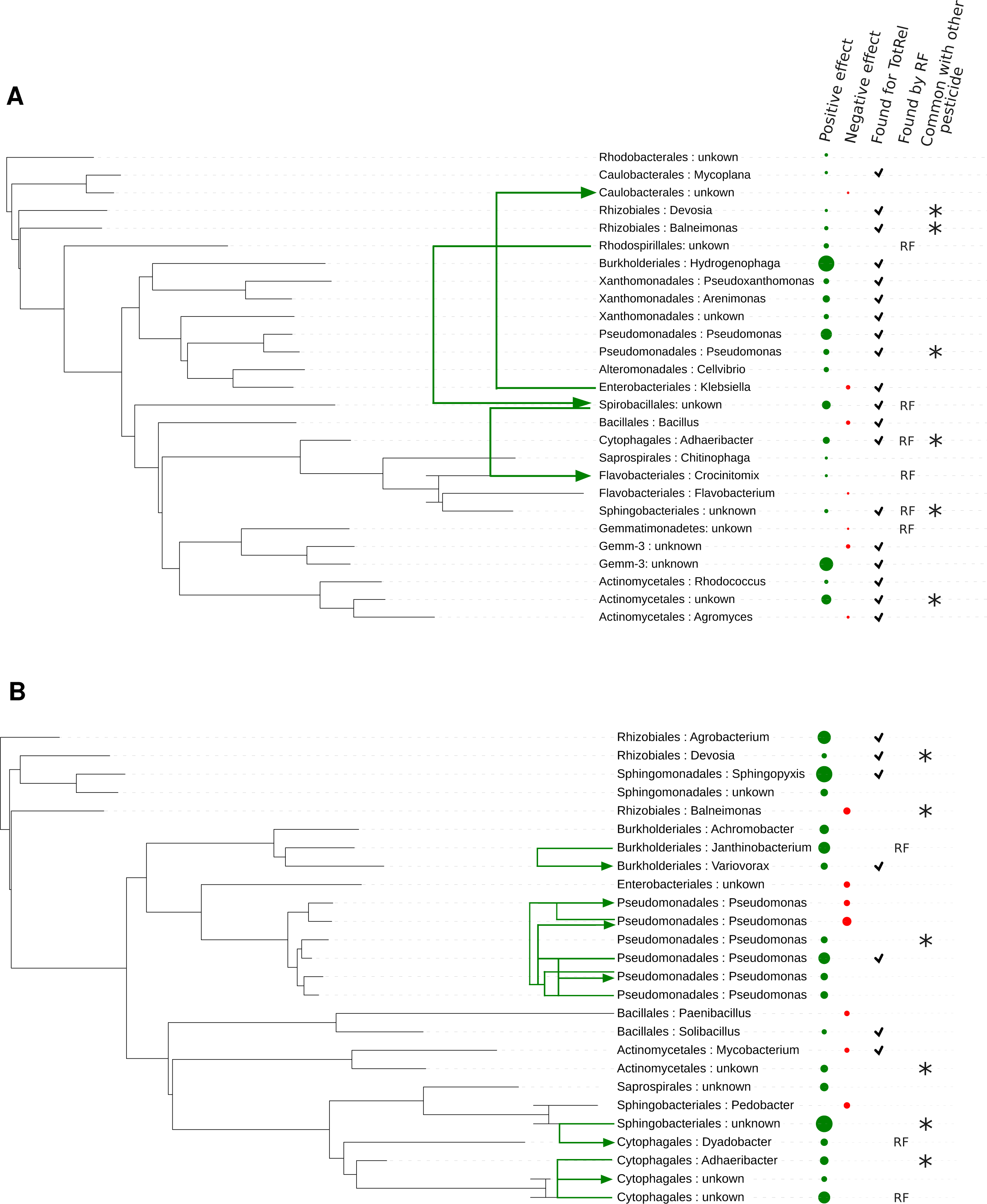
Taxonomic classification, importance and relationship between OTUs with highest predicted effect. Taxonomic classification (”order: genus” if known (otherwise “unknown”)) of OTUs having highest estimators predicted by BL and BRR for the Slope phenotype for A) glyphosate and B) isoproturon. Green arrows link OTUs with significant co-occurrence pattern, determined by positive partial correlation. Dot size displays the effect size of positive estimates (green) and negative estimates (red). Checks indicate OTUs also detected among the most important for the phenotype Total by BL and BRR, then “RF” column indicates OTUs detected by RF method on Slope phenotype. Finally, stars denote OTUs that are detected for both GLY and IPU.

### Mixing compositional variants and evaluating the GLY and IPU mineralisation potential of the resulting communities

The next step consisted in mixing GLY with IPU compositional variants in order to construct multidegrading communities and inoculate them into the non-degrading sterile Ecotron soil. We tested six different mixing scenarios in order to detect which community characteristics could impact the degrading ability of resulting communities. Interestingly, only slight differences were found between the six tested scenarios when comparing the GLY and IPU mineralisation potential of the resulting communities compared to their parental ones (Fig. 5). These results indicate that no strong antagonism between bacterial populations associated with the degradation of IPU and GLY exists, but also that no overyielding of the mineralisation potential was detected. However, we observed that the choice of the phylogenetic diversity level of the parental communities impacted the mineralisation potential of the resulting communities. On the one hand, choosing parental IPU communities with low phylogenetic diversity led on average to lower IPU mineralisation potential in the resulting communities. On the other hand, choosing parental GLY communities with high phylogenetic diversity resulted on average in communities with higher GLY mineralisation potential (Fig. 5).

**Figure 5:**
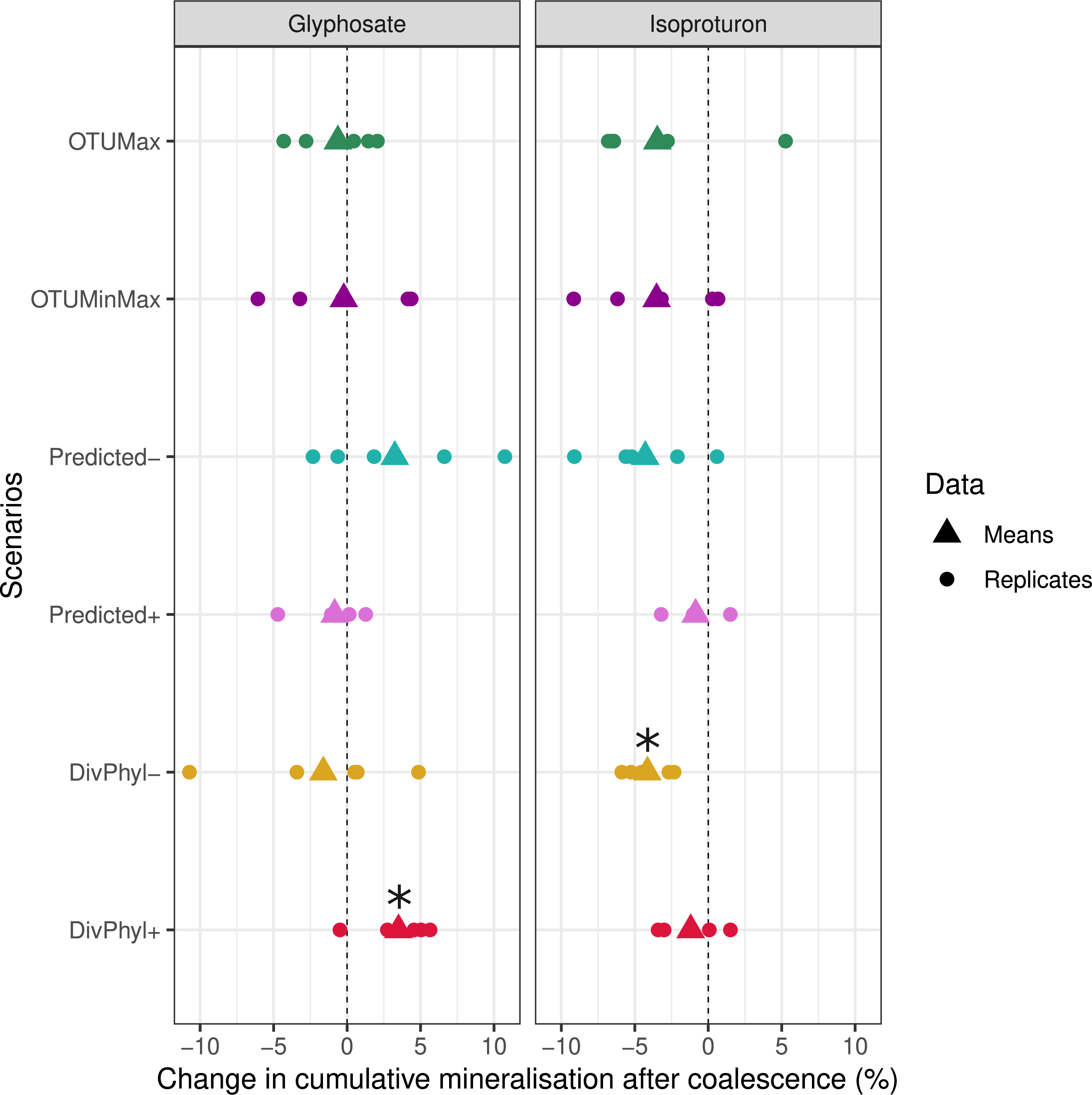
Cumulative mineralisation differences between original and coalesced communities depending on original communities’ properties. Mineralisation difference of total cumulative percent after 42 days, with n = 5 for each scenario to choose coalescent communities based on their properties. Stars show distributions with mean significantly different from 0. The two first scenarios depend on OTU estimated effects on pesticide mineralisation, first maximizing the abundance of OTUs with a positive effect while minimizing the abundance of OTUs with a negative effect, second maximizing OTUs with positive effect times their abundance. Other scenarios result from trade-off between measured pesticide mineralisation and either prediction accuracy on other pesticide or community diversity (phylogenetic difference).

### Coalescence of the herbicide multi-degrading communities into a non-degrading soil

Finally, we tested the efficiency of coalescing the resulting multi-degrading communities into the non-degrading non-sterile Ecotron soil by selecting the five best communities. We evaluated different inoculation ratios and results are contrasted, depending on the herbicide (Fig. 6). For GLY, non-inoculated control showed relatively high mineralisation ability averaging 21.7% after 42 days of incubation, attesting the mineralisation ability of the indigenous microbial community. Furthermore, low inoculation ratios did not increase this potential and high inoculation ratios (5 g mixed in 50 g) produced only a slight increase in GLY mineralisation ability (mean of 24.9%). On the other hand, IPU mineralisation potential was quite low in the non-inoculated control (mean of 11.1%). It increased significantly and strongly in response to inoculation of even small quantities of multi-degrading communities (mean of 28.4% for 0.1 g, going up to 42% for 1 g).

**Figure 6:**
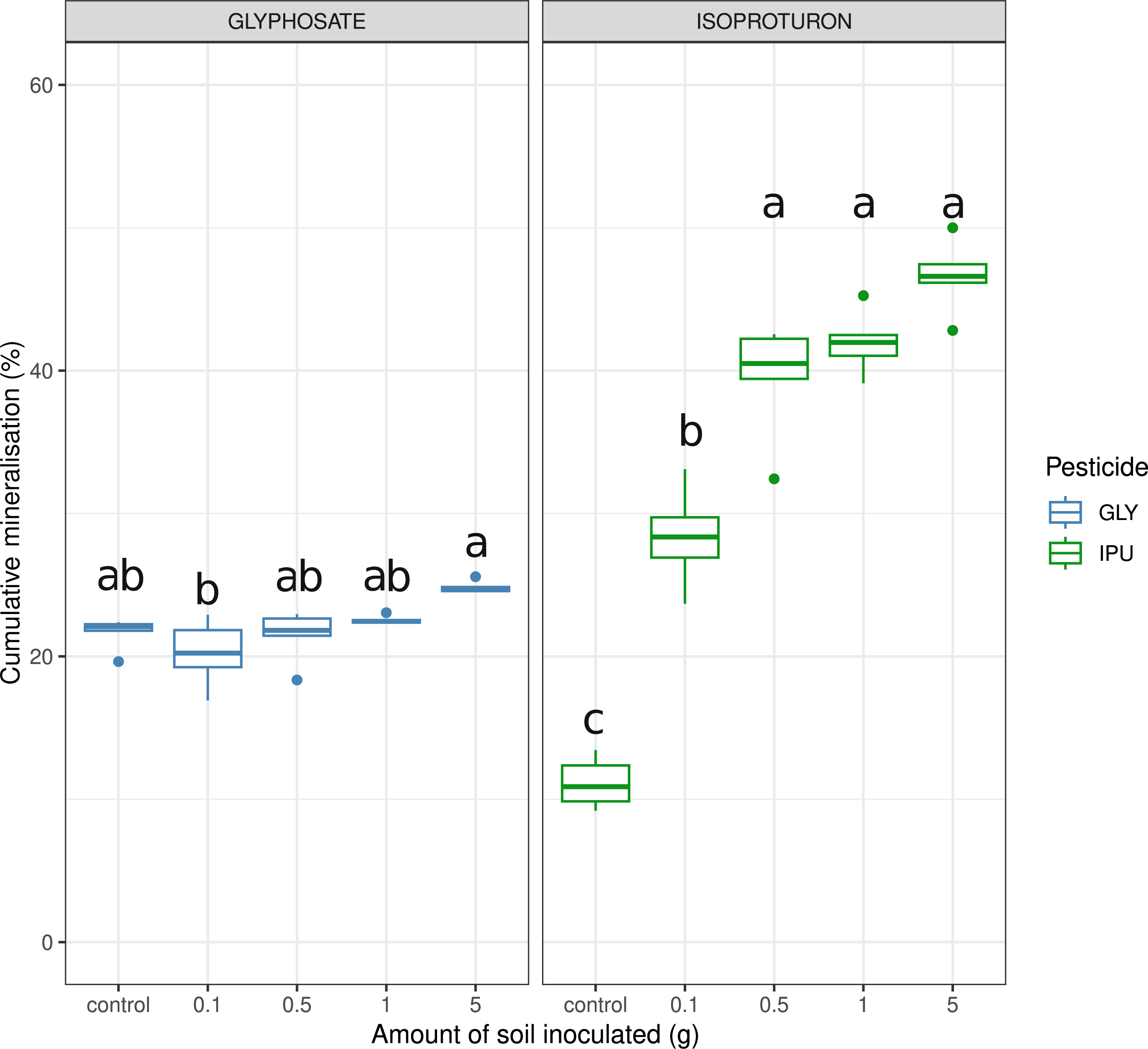
Cumulative mineralisation of GLY and IPU in pesticide spiked soil microcosms inoculated with multidegrading community. For each gradual inoculation n = 5, from 0 (control) to 5 g of soil in 30 g of non-sterile soil. Cumulative mineralisation is showed after 42 days of incubation.

**Figure 7:**
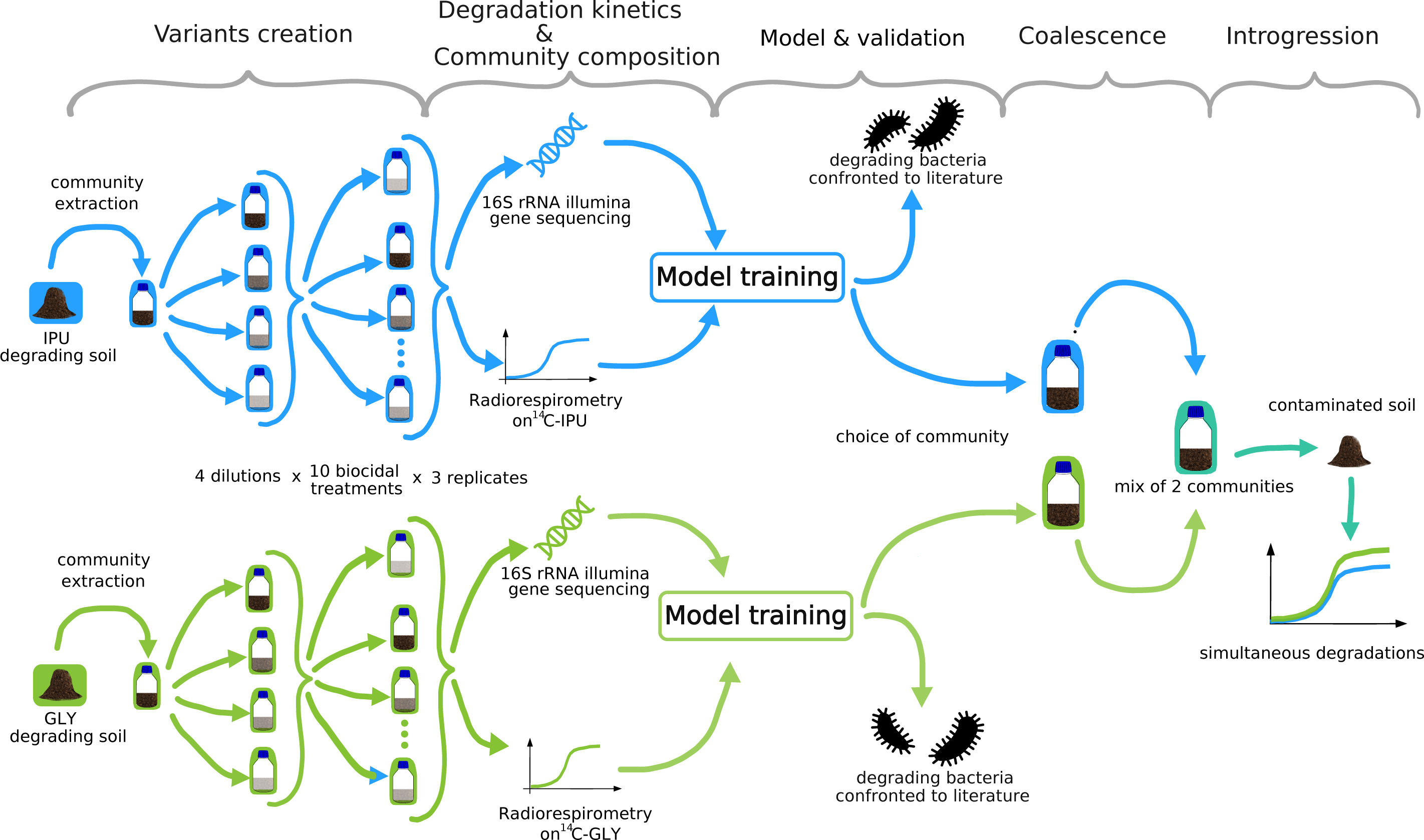
General layout of the experimental strategy.

## Discussion

Environmental filtering refers to the process by which local abiotic conditions select organisms that are able to cope with the existing physico-chemical properties (14). It is thought to be one of the main processes causing the inefficiency of *in situ* microbiome engineering (15; 35; 36). Here, we chose to *γ*sterilize the target environment, *i. e.* the environment in which bioremediation of a pollutant is targeted, and use it as a first screening for degrading microbial communities. After community extraction, manipulation and inoculation of the target sterilized soil, we observed that herbicide mineralisation potential was kept at high levels and even exceeded the original soils mineralisation potential in the case of IPU (Fig. 2). The bacterial community structure of the manipulated GLY and IPU mineralising communities was drastically altered compared to the original soils from which they originated (Supp. Fig. 3), probably in relation with the sandier structure and the more acidic conditions of the target environment. This confirms the strong role of environmental filtering in shaping microbial communities, albeit without impairment of their functioning in our case.

We combined serial dilution to biocidal treatments to create compositional variants of GLY and IPU mineralising bacterial communities. Consistent with theory and previous studies (14; 37), dilution had a strong impact on bacterial richness (Fig. 2), higher levels of dilution resulting in less diverse communities illustrating the loss of rare taxa. In the case of GLY, this decline in bacterial richness was associated with a decrease in the mineralisation capacity, underlining a relatively straightforward link between diversity and functioning often reported in the literature (38; 39; 40). However, in the case of IPU degrading communities, we did not observe such a relationship, suggesting that the IPU mineralisation function might be supported by a specific group of originally common OTUs, which might even have benefited either from the community manipulation or the target environment local conditions we subjected them to. This interpretation is supported by the general knowledge that IPU mineralization abilities is shared only among a small amount of bacteria genus (41).

The successful implementation of our prediction approach relied on our ability to generate a compositional continuum of communities varying in their capacity to degrade the pesticides. A strong level of structure in the composition of the communities following our treatments could be detrimental to the overall ability of the model to yield satisfactory predictions. The same restriction is faced in genomic prediction and regular updates with new data (phenotype-genotype information) are carried out to keep the model adjusted over time (42). Our success in obtaining a continuous distribution of community variants notwithstanding, the transfer of genomic prediction tools on microbial function is challenged by two main differences in data: presence/absence of alleles are replaced by a more continuous and potentially limitless count of OTUs, and the likely change in community composition over time compared to the fixed genetic information. This last point turned out to be particularly relevant in IPU case, with clearly superior predictions for starting mineralisation rate than final mineralisation potential (Fig. 3). For future consideration, monitoring the change of degrading communities over time seems crucial for our ability to devise prediction models trained on data that are not too divorced from the data they try to predict. Despite all these potential pitfalls, our approach gave very satisfying preliminary results and is supported by promising results from machine learning methods tested on microbial communities to predict community function (43; 44).

There was no clear distinction in prediction accuracy between the three statistical methods that we used (BL, BRR and RF). In parallel, and to get an idea of the intrinsic properties of microbial communities that could influence the statistical power of these methods, we used simulations of degrading capacities of an existing OTUs table postulating various “trait architecture” (*i. e* various ways for OTUs to contribute or not to the simulated phenotype). The results showed that predictions were better when the simulated phenotype was underlain by a few abundant OTUs (Supp. Fig. 5). This type of architecture matches well what we observe for IPU degradation, which seems to be performed by a few specific OTUs (45), and consistent with the fact that our model gives a very high prediction accuracy. The case of GLY is different, with the degrading potential seemingly more spread out across OTUs, yet we obtained reasonably accurate predictions there too.

To validate the model predictions, we looked at OTUs with the highest and lowest predicted effect sizes and checked whether they were already identified in previous literature. Two *Pseudomonas spp.* strains are identified for their GLY degrading capacities, which matches 2 OTUs with positive estimates in our analysis (46; 47). On the opposite, two OTUs with negative estimate, *Bacillus cereus* and *Flavobacterium sp.*, show isolated strains for their GLY degrading capacities (46; 47). These last apparently mixed results could be explained by a competition pattern related to phylogeny, pattern well known in microbial ecology (48). Indeed, we noticed that phylogenetically close OTUs often turned out to have opposite effect on degradation function, suggesting that they might be in competition, with one bearing the degrading capacity and the other occupying a similar ecological niche. These non-degrading competitors would expand in absence of degrading OTUs, and therefore be negatively correlated with pesticide mineralisation.

Concerning IPU, degrading strains are mostly found within the *Sphingomonas* genus (49; 50), which matches our model prediction with two OTUs with positive estimates belonging to this taxon. The other OTUs with high prediction power have not been yet described as IPU degraders but are good candidates either directly involved in the degradation or facilitated by degrading OTUs.

Increasing interest for community coalescence as a widespread microbial ecological event and recent studies (51; 52) prompted us to establish coalescence scenarios based on community-level properties. By doing so, our initial objective of building multi-degrading communities turned out to be a great success. All coalesced communities proved to be efficient at degrading both pesticides. However, among all tested community properties, only the phylogenetic diversity was revealed as an important community feature to maintain high degradation capacities for both pesticides after coalescence.

Moreover, we can hypothesize from our results that no strong antagonisms exist between IPU and GLY degrading community members, which is consistent with the chemical structure discrepancy between the two pesticides and their degrading pathways (45; 46). This suggest a modular coalescence in which input communities are preserved and only limited new interactions between communities are established, as opposed to chimeric coalescence showing numerous new interactions (51).

Introgression of the multi-degrading communities in an inhabited polluted soil resulted in contrasted outputs between the 2 pesticides. A clear transfer of IPU degradation capacity was observed, even at low inoculation levels. This success can be explained by the establishment of cohesive unit of co-selected taxa along the experiments for their use of IPU as substrate, a hypothesis supported by previous theoretical work (53). Conversely, introgression of GLY degrading capacities gave mixed results, consistent with an already present degrading potential. Indigenous community benefits of the priority effect (54), making it difficult for new taxa to establish. At high inoculation level though, degradation is enhanced, either demonstrating the importance of invasive pressure (55; 15), or the additive relationship between species richness and community phenotype (40; 52).

## Conclusion

Microbiome engineering is a fast-developing research field. It is a promising approach for bioremediation purposes if inoculants are able to effectively establish and fulfill the functions they were selected for in contaminated soils. We propose here an innovative and efficient strategy for building multidegrading bacterial communities that not only address the challenges posed by the environmental factors, but also considers biotic interactions between community members. Using a coalescence framework, our approach allowed to efficiently establish and perform the desired functions in the target soil.

## Acknowledgements

We thank members of the EMFEED team for their help and support, and specifically Laurent Philippot for his inputs. This work was partially supported by grants from the INRAE HoloFlux Metaprogramme (INT-BXL project).

## Author Contributions

ST, MS & AS designed the study; ST, MD-L, SR & NR performed the experiments; ST, JA, SR, MS & AS analyzed the data; ST wrote the manuscript; MD-L, FM-L, SR, MS & AS edited the manuscript.

## Data availability statement

All data and codes used during this study have been deposited to Zenodo. Raw sequences were deposited at the NCBI under the BioProject PRJNA1075697.

