## Supplementary material for "Engineering multi-degrading bacterial communities to bioremediate soils contaminated with pesticides residues": Table 1

Table 1: Biocidal treatments applied on herbicide-degrading communities

| Treatment | Abbreviation | Modality |
| --- | --- | --- |
| Ciprofloxacin (antibiotic) | Cipro | 5h incubation at 25°C with  ciprofloxacin at 66 *µ*g/ml |
| Ramoplanin (antibiotic) | Ramo | 5h incubation at 25°C with ramoplanin at 70 *µ*g/ml |
| pH2 | pH2 | 2h incubation at 25°C with 730*µ*l acid malic 1M then 3 washes |
| pH11 | pH11 | 2h incubation at 25°C with 370 *µ*l ammoniac 20% 1M then 3 washes |
| Oxidant 1 – strong | Ox1 | 1h incubation at 25°C with 730*µ*l  H202 0,98M |
| Oxidant 2 – mild | Ox2 | 1h incubation at 25°C with 295*µ*l  H202 0,98M |
| Heat shock | HS | 0°C for 5min ; 70°C for 15min; 0°C for 5min |
| UV | UV | 2h exposition to UVC light |
| Freeze-thaw | FT | 6 x (-80°C for 15min ; 30°C for  15min) |
| Control | CT | / |
