## Supplementary file for "Engineering multi-degrading bacterial communities to bioremediate soils contaminated with pesticides residues"

### Supplementary Informations

#### Supplementary figure 1

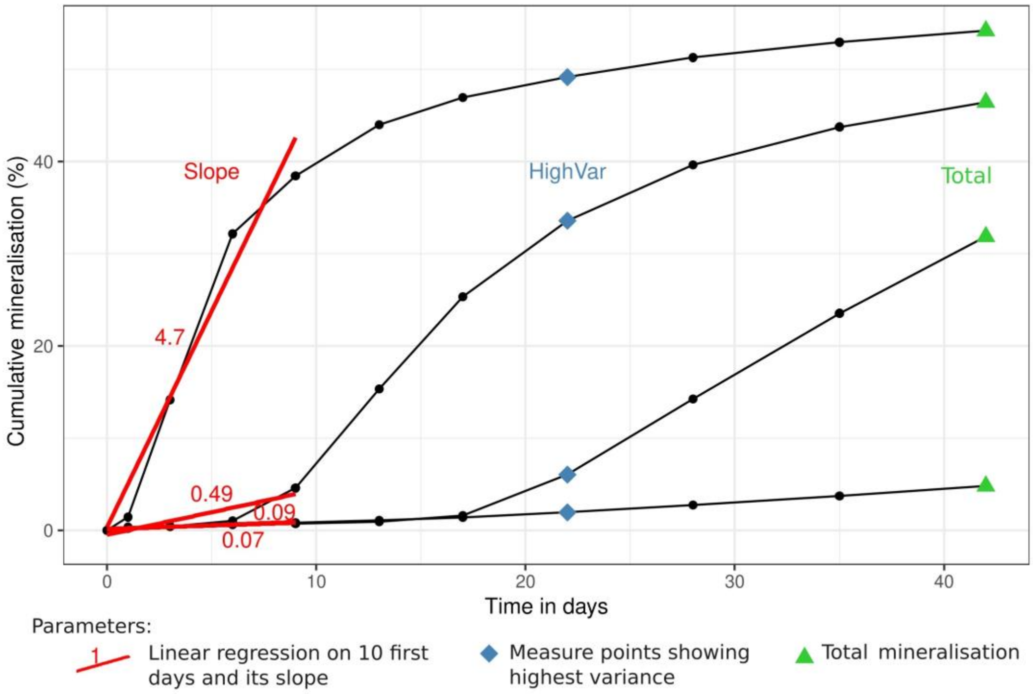


Supp. Figure 1: **Measured parameters on mineralisation curve.** These 3 parameters are to be predicted by statistical methods: Slope (slope of linear regression on the 10 first days), HighVar (mineralisation value at the time point at which the variance across samples of the cumulative mineralisation potential is maximised) and Total (total relative mineralisation on last day)

#### Supplementary figure 2

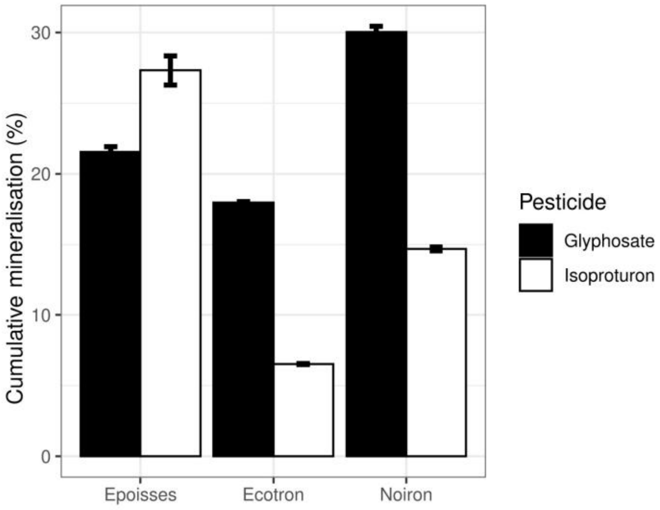


Supp. Figure 2: Mineralisation potential of GLY and IPU of the three soils used in this experiment. Mineralisation was measured by radiorespirometry (22 days) using 14Cphenyl ring labelled IPU and 14C-GLY. Cumulative percentage of 14CO2 evolved from 14C pesticide mineralisation after 22 is indicated. Error bars represent standard error of the mean (n=2).

#### Supplementary figure 3

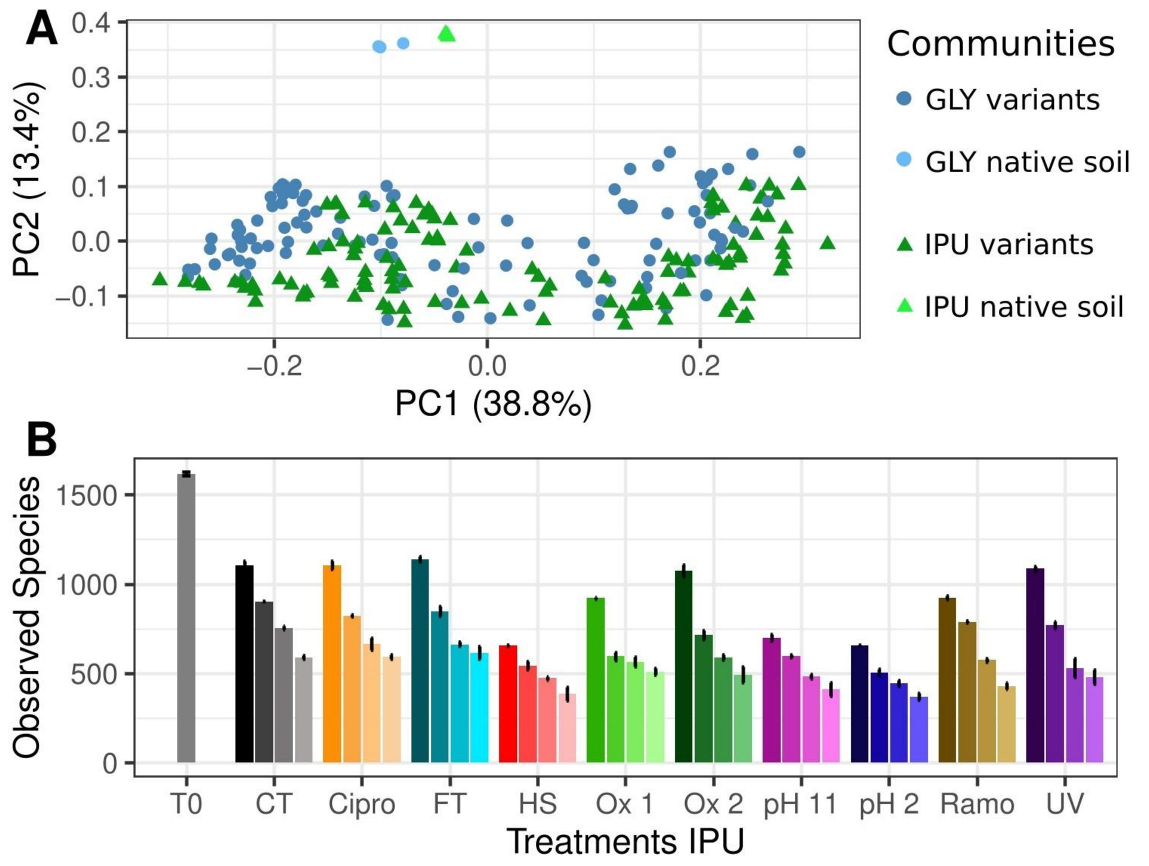


Supp. Figure 3: **Bacterial diversity of the native and variants communities**. A. Principal Coordinate Analysis performed on weighted UniFrac distance measured between communities. B. Number of observed species (OTUs) displayed for each treatment coupled with dilution. For each bar is the mean of 6 communities (3 IPU-degrading communities and 3 GLY-degrading ones, error bars representing standard error of the mean.

#### Supplementary figure 4

From the three statistical methods we ran on the Slope phenotype, we looked further in the OTUs detected has having a large impact on the phenotype. It is a way to test the explanatory power of the methods, and to compare them between each other. Strikingly, methods resulting OTUs are shared for BL and BRR method while OTUs detected by RF are quite different (Fig. 11). This can be a reflect of the closeness of the two linear methods (BL and BRR), but also a logical outcome of the intrinsic non linear functioning of the RF, and the lack of explanatory power associated (Fig. 12). We therefore focused more precisely at the taxonomy and the partial correlation patterns between OTUs detected by both BL and BRR models only.

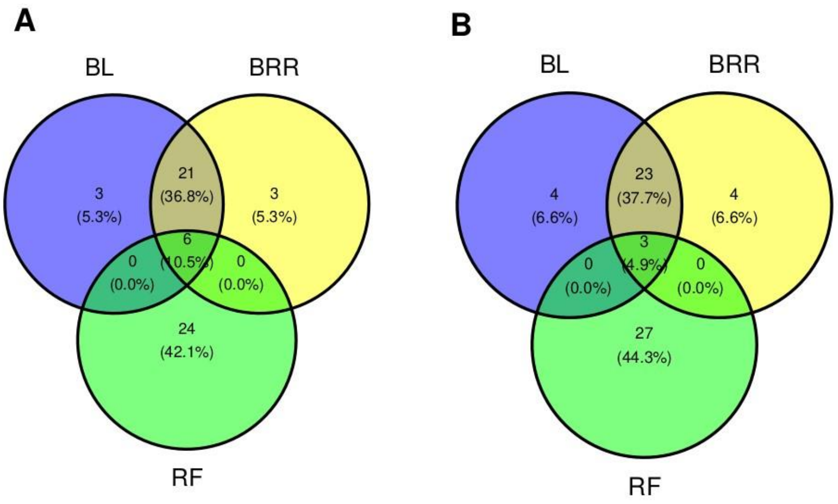


Supp. Figure 4: **Venn diagrams on the 30 OTUs with highest effect detected by BL, BRR and RF for Slope phenotype**. Display of OTUs detected by the 3 statistical methods and how many are in common for A. GLY prediction and B. IPU prediction.

#### Simulations

**Materials & Methods** With the aim of evaluating the prediction power of the three statistical approaches, namely Random Forest (RF), Bayesian Ridge Regression (BRR) and Least Absolute Shrinkage and Selection Operator (LASSO), we conducted analyses on simulated data. We linearly associated composition of bacterial communities (from an existing data set) to a trait, or function, hereafter denoted as phenotype. The data set is composed of OTUs from 256 communities sampled across a single field in INRAE's Époisses experimental domain in 2020. The data set had been previously filtered to get rid of very rare OTUs, with a threshold of 250 counts, resulting in about 1000 OTUs in total. Looking at the data, we added a filter on the communities and got rid of the ones with very low number of reads (below 2000 reads), for a mean of 5641 reads on all the communities (and a standard deviation of 670). After visualisation of data and preliminary prediction results, we noticed bias due to extreme OTUs distributions. A few OTUs showed very large counts in one or two communities and very low (around zeros) in others. Such distributions probably usually arose from technical abnormalities. These particular OTUs were taken out of the analysis through a purposely developed criterion: ration of OTU distribution variance on variance of this same distribution without the maximum value. We then empirically set a threshold of 4.5 looking by hand at the different OTUs distributions. Finally, the counts were scaled to account for differences in sequencing depth between communities by dividing them by the total read depth by sample and multiplying these by the maximum read depth found. Four scenario of phenotype construction were tested : • Scenario 1: 1 % of all OTUs contribute in the phenotype and those are the more abundant ones (considering the abundance as the sum of all counts across communities for any given OTU) • Scenario 2: 1 % of all OTUs contribute in the phenotype and those are randomly chosen among the 99 % of the less abundant • Scenario 3: 10 % of all OTUs contribute in the phenotype and those are the more abundant ones • Scenario 4: 10 % of all OTUs contribute in the phenotype and those are randomly chosen among the 90 % of the less abundant

For each of the contributing OTUs, an effect on the phenotype was randomly drawn from a Normal distribution with mean 0 and standard deviation of h2 divided by the number of participating OTUs, with h2 included in [0, 1]. h2 is directly equivalent to the heritability in genetics and accounting for the proportion of phenotypic variance due to the community composition vs the part influenced by the environment. It was set at 0.5 for the rest of the analysis. The phenotype was then computed as the sum of the counts of participating OTUs times their effect plus an error term randomly drawn from a Normal distribution with mean 0 and standard deviation of 1-h2. Finally, prediction models were used to link simulated phenotype to the communities data (OTU table) with 5-fold CV. Linear models BL and BRR were run for 5 000 iterations with a burn-in of 1 000 iterations. The resulting effects are calculated from mean estimated parameters ($\lambda$ for BL and $\mu$, $\sigma$ for BRR) and the RF was run for 500 trees with node size 5. We used two metrics to assess the performances of the models: correlation between predicted and realized phenotype, reflecting the prediction power, and the match between OTUs with large predicting effect and the ones that effectively participated in the simulated phenotype as a way to measure explanatory pertinence.

**Results** Results reveal clear differences between linear models (BL and BRR) and Random Forest, depending on scenarios (Fig. 12 B). RF correlation between predicted and simulated phenotype is close to 1 (comprise between 0.98 and 0.99) for every scenario while linear model prediction is largely lower for scenario based on rare OTUs : for scenario 2, BL gives a correlation around 0.85 +- 0.13, BRR 0.76 +- 0.13 and for scenario 4, BL is at 0.50 +- 0.1 and BRR 0.64 +- 0.08 compared to higher numbers for the other scenarios : 0.99 for both models in scenario 1 and for scenario 3 0.95 +- 0.01 with BL and 0.94 +- 0.02 with BRR. Larger variability is observed for the lower results, for different runs but same simulation (black error bars) and between simulations (change of OTUs participating to the phenotype). Then linear models are sensible to simulated phenotype construction with a loss of predictive power and replicability when the phenotype is based on rare OTUs, compared to abundant OTUs.

The success score also shows the divergence between rare OTUs scenarios (2 and 4) and abundant one (1 and 3), with very low score for all 3 methods in the first case (between 10% and 30%) (Fig. 12 A).

For the two scenarios based on abundant OTUs however, linear models gave higher success rate than RF, without a clear difference between BL and BRR (in scenario 1, score of 85%+-12 for BL, 74%+-21 for BRR while RF is at 40%+-13 and for scenario 3: 93%+-6 for BL, 96%+-8 and 63%+- 21 for RF).

Scores are higher for scenario 3 surely due to the higher number of contributing OTUs (from 10 to 100).

We see that statistical methods performances depend greatly on OTUs linearly participating in simulated phenotypes (or trait architecture), with best results obtained when only abundant OTUs are at stake. This result turns out to be supportive for our experimental design, since we hypothesise that advantages of herbicide degrading OTUs would lead them to be numerous relatively to others. The second highlight lies in the difference between RF and linear models, the first one showing good prediction capacities but poor explanatory power. On the other hand, linear models gave good predictions in favorable scenarios and satisfactory explanatory results (scores) in the same context.

This discrepancy can be used to determine the closest scenario on experimental data: if linear models laid significantly lower and highly variable results, rare OTUs are at stake for the phenotype.

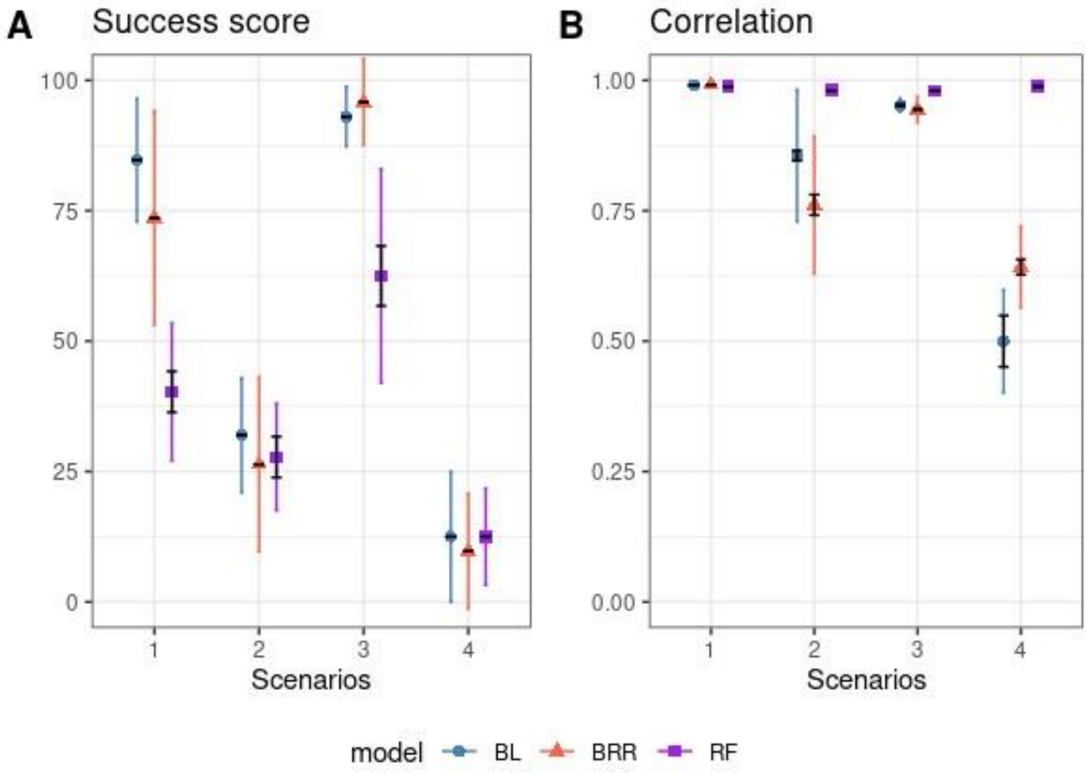


Supp. Figure 5: **Sensitivity analysis of prediction methods performance.** Black error bars represent standard error of the mean for runs with the same simulated phenotype and participating OTUs with n=8 (intrinsic variations). Colored errors bars are standard deviation of the mean for phenotype simulation runs with n=8. (A) Success score defined as percentage of OTUs in the top 10 largest estimates actually contributing to the phenotype. (B) Correlation between predicted and realized phenotype values
